# Seasonal stability and dynamics of DNA methylation in plants in a natural environment

**DOI:** 10.1101/589465

**Authors:** Tasuku Ito, Haruki Nishio, Yoshiaki Tarutani, Naoko Emura, Mie N. Honjo, Atsushi Toyoda, Asao Fujiyama, Tetsuji Kakutani, Hiroshi Kudoh

## Abstract

Organisms survive in naturally fluctuating environments by responding to long-term signals, such as seasonality, by filtering out short-term noise. DNA methylation has been considered a stable epigenetic mark but has also been reported to change in response to experimental manipulations of biotic and abiotic factors. However, it is unclear how they behave in natural environments. Here, we analyzed seasonal patterns of genome-wide DNA methylation at a single-base resolution using a single clone from a natural population of the perennial *Arabidopsis halleri*. The genome-wide pattern of DNA methylation was primarily stable, and most of the repetitive regions were methylated across the year. Although the proportion was small, we detected seasonally methylated cytosines (SeMCs) in the genome. SeMCs in the different contexts showed distinct seasonal patterns of methylation. SeMCs in CHH context were detected predominantly at repetitive sequences in intergenic regions. Additionally, we found that CHH methylation within *AhgFLC* locus showed a seasonal pattern that was negatively associated with changes in gene expression. Gene-body CG methylation (gbM) itself was generally stable across seasons, but the levels of gbM were positively associated with seasonal stability of RNA expression of the genes. These results suggest the existence of two distinct aspects of DNA methylation in natural environments: sources of epigenetic variation and epigenetic marks for stable gene expression.

## Introduction

DNA methylation at cytosine residues is an epigenetic mark that can be maintained through cell divisions in a wide range of eukaryotic genomes (1).The previous analyses in diverse organisms have revealed genomic distributions of DNA methylation vary among organisms (2-7). In plants, DNA-methylation varies both between and within species (8), and sometimes is associated to phenotypic variation (9-11). Although the level and patterns of DNA methylation are heritable to a certain extent, the mechanisms that produce and maintain epigenetic variation across generations are largely unknown.

DNA methylation can vary between individuals also by non-genetic causes. It has been shown that both biotic and abiotic treatment can modify DNA methylation (12-14). Because of its semi-stable and semi-labile nature, non-genetic changes in DNA methylation are not explained by simple environmental effects. For example, even in a genetically homogeneous background under stable laboratory conditions, epigenetic variation in DNA methylation can occur during repeated self-pollination in a transgenerational manner (15, 16). Therefore, it is difficult to predict whether DNA methylation is stable or dynamic under natural conditions. Recently the need for ‘*in natura*’ studies has been highlighted in order to understand how organisms respond to environmental signals by filtering out innumerable fluctuations and noise (17, 18). In the temperate regions, seasonality is the most prominent cause of environment fluctuations. However, we still do not understand how DNA methylation behaves across seasons.

In order to reveal the seasonal dynamics of DNA methylation, here, we conducted an ‘*in natura*’ study on genome-wide DNA methylation using a single clone growing in a population of *Arabidopsis halleri* (L.) O’Kane & Al-Shehbaz subsp. *gemmifera* (Matsum.) O’Kane & Al-Shehbaz (hereafter referred to as *A. haleri*), a close relative of *Arabidopsis thaliana*. Its clonal propagation and perennial life-cycle allowed us to sample leaves all year round from the single clonal individual (19). We studied seasonal dynamics of key flowering-time genes and whole transcriptome previously in the site (20, 21). In this study, to understand the dynamics of DNA methylation, we performed whole-genome bisulfite sequencing (WGBS) at 1.5-month intervals, over a year, under natural conditions. We adopted the strategy of monitoring the seasonal pattern in a single clonal individual with a uniform genetic background.

DNA methylation occurs in three contexts, according to the flanking sequence, i.e. CG, CHG and CHH (H = A, C, or T). The former two form symmetrically and the latter one asymmetrically, in terms of sequences on the complementary strands. Since these contexts of methylation are distinctly regulated and associated with DNA replication, histone modification, and non-coding RNA production (22-24), we examined seasonal patterns of DNA methylation by conducting single-base resolution analyses.

The results suggested the existence of seasonally methylated cytosines (SeMCs) in the genome, at least for the examined clone. Interestingly, DNA methylation changes occurred in a context-dependent manner. There were distinct patterns of seasonal changes among CG, CHG, and CHH methylation. Moreover, our analysis revealed that genic CG methylation, i.e. gene-body methylation (gbM), was seasonally stable by itself, and was associated with seasonal stability of RNA expression. This study suggested not only that there is a dynamic nature in DNA methylation in plants in their natural habitats, but also highlights the implications of DNA methylation in robust maintenance of stable RNA expression.

## Results

### Seasonal stability in large-scale distribution of DNA methylation

To investigate the dynamics of DNA methylation in a natural environment, we analyzed genome-wide DNA methylation in a natural population of *A. halleri* (Fig. 1 *A* and *B*). In the study site, the hourly air temperature ranged from −4.3 °C to 36.3 °C during the one-year study period, from Nov 2014 to Sep 2015 (Fig. 1*C*). We collected leaves from a single clonal patch of *A. halleri* at 8 sampling times, at 1.5-month intervals across a year, and performed WGBS (Fig. 1*C*). From this, we obtained a series of genome-wide DNA methylation data (Table S1).

**Fig. 1.**
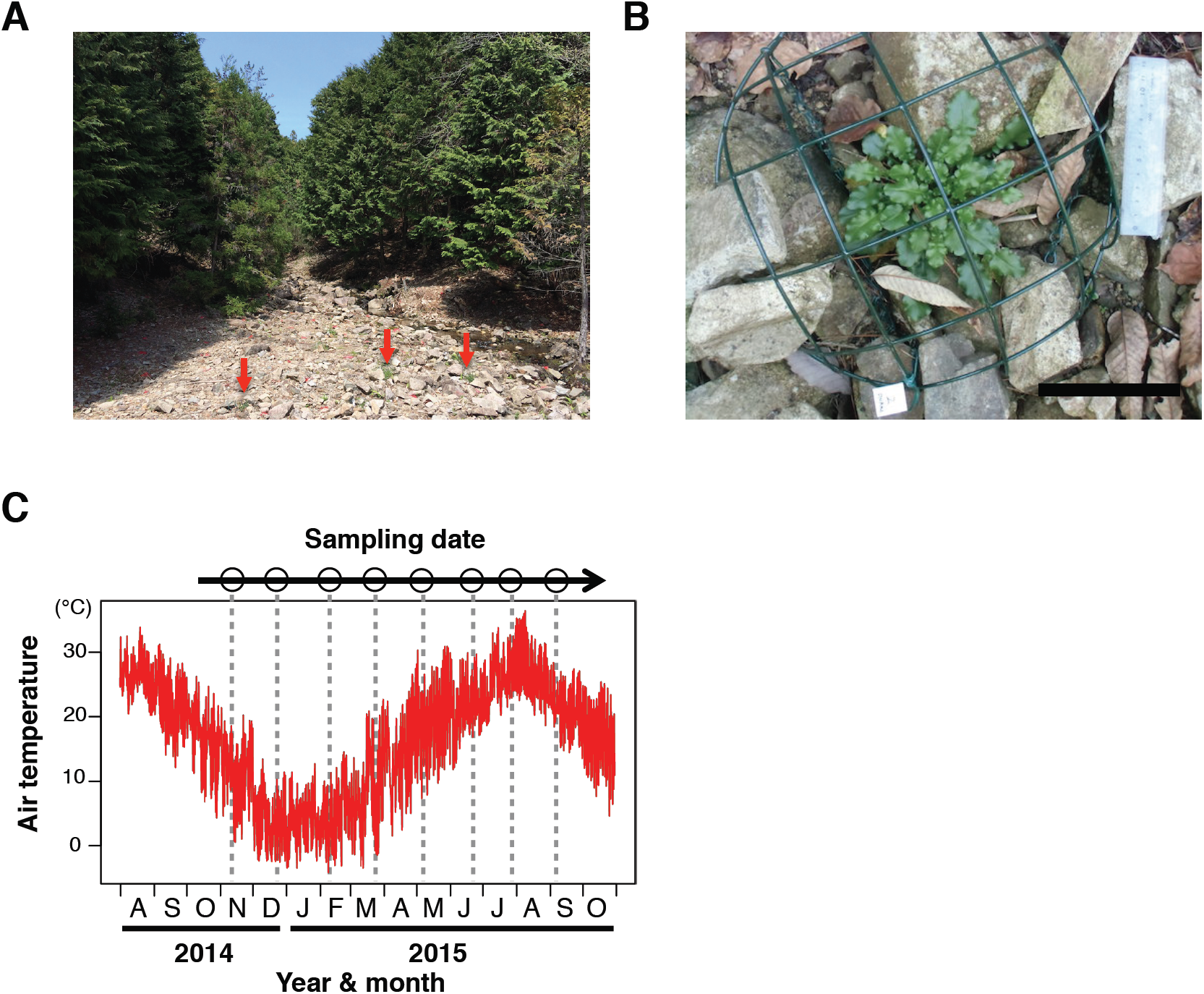
Sampling site and dates for the seasonal DNA-methylation analysis in a natural population of *Arabidopsis halleri* subsp. *gemmifera.* (*A*) A photograph of the study site alongside a small stream through the *Cryptomeria* and *Chamaecyparis* plantation in Hyogo Prefecture, Japan. Red arrows indicate individuals of *A. halleri.* (*B*) An individual of *A. halleri* in the study site. The cage was used for protection against deer herbivory. White bar indicates 100 mm. (*C*) Sampling dates (8 time points) and temperature regimes during the study period. Sampling was performed every ca. 1.5 month for a year. The red line indicates hourly air temperature, and dotted lines represent timings of the sampling.

Bulk DNA methylation levels were relatively constant across the eight time points, and ca 45%, 20%, and 6% of cytosines in the genome were kept methylated in CG, CHG, and CHH contexts, respectively (Fig. S1*A*). The large-scale distribution of DNA methylation was determined by the positions in the genome, as represented by radial patterns in the circos plot for the longest 30 scaffolds (Fig. 2*A*). The distribution of methylated sites remained constant across the 8 sampling times (represented by 8 concentric circles for each methylation context in Fig 2*A*). We observed conspicuous aggregation of DNA methylation on repetitive sequences (Fig 2*A*). For example, scaffolds 3 and 18 is characterized by low and high density of repetitive sequences that corresponded to relatively low and high methylation levels, respectively (Fig 2*A*). In the comparative analysis between genic and repetitive regions for the whole genome, the level of DNA methylation in repetitive sequences was higher than that in genes (Fig. S1*B*). A similar pattern was confirmed by an analysis using 100 kbp windows for the whole genome (Fig. 2*B*). The level of DNA methylation was correlated with density of repeats in each window for all three contexts in all samples (e.g. Fig. 2*B* for Nov. 2014, Pearson’s correlation coefficients were 0.64, 0.68, and 0.70 for CG, CHG, and CHH context, respectively; Table S2 for the other sampling times). Although aggregation of DNA methylation at repetitive sequences was similar to the patterns previously reported in related Brassicaceae (2, 3, 25), seasonal stability in large-scale distribution of DNA methylation was reported for the first time here.

**Fig. 2.**
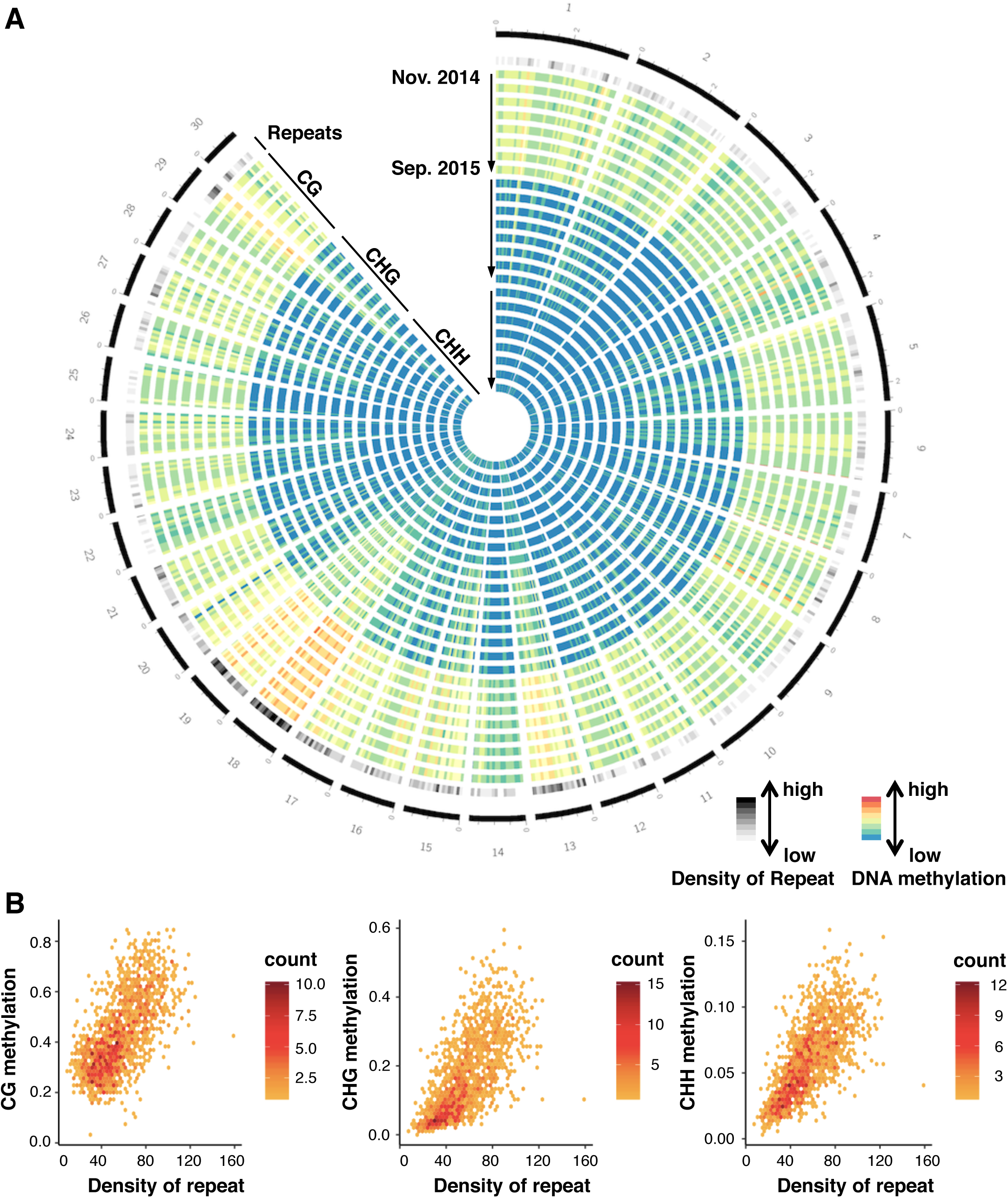
Genomic pattern of DNA methylation across a year. (*A*) A circos plot showing seasonal patterns (8 time points) of DNA methylation in CG, CHG, and CHH contexts for the longest 30 scaffolds of *A. hallleri.* On the outermost circle, scaffold positions are indicated by numbers and black bars with scales (one scale = 0.4 Mbp). Each shaded/colored circle represents the scaffold-wide distribution of genomic attributes for each 100-Kb window. The second outermost circle represents density of repetitive sequences (including transposable elements). Each tile indicates the density: the lowest in white, the highest in black (0–0.5). The next 24 circles represent DNA methylation levels at 8 time points (starting from Nov 24, 2014 to Sep 8, 2015, towards the inner circles shown by the arrows) for CG, CHG, and CHH contexts, respectively. Colors in each tile indicates level of methylation: the lowest as blue, the highest as red (0–0.9 for CG context, 0–0.5 for the others). (*B*) Scattering plots comparing CG, CHG, and CHH DNA methylation against TE density for all 100 kbp windows across the 30 scaffolds.

### SeMCs: Seasonal dynamics of DNA methylation

Next, we searched for SeMCs by detecting differently methylated cytosines in the genome across the eight time points (Fisher’s exact test; *P* < 0.001; FDR < 0.2). The majority of cytosines did not show statistical differences in methylation across the year, and the proportion of SeMCs was less than 0.5% of total cytosines (Fig. S2 *A* and *B*). Still, we detected 62,716, 47,140, and 179,385 SeMCs in CG, CHG, and CHH contexts, respectively (Fig. 3*A*). They showed diverse seasonal patterns in their level of methylation across the year (Fig. 3*A*). Interestingly, each context of DNA methylation showed distinct patterns of seasonal change: the number of SeMCs that peaked in a particular month was the highest in July, March, and September in CG, CHG, and CHH contexts, respectively (Fig. 3B). Differences among contexts of SeMCs would reflect the responsiveness of the regulatory mechanisms of DNA methylation to environmental cues.

**Fig. 3.**
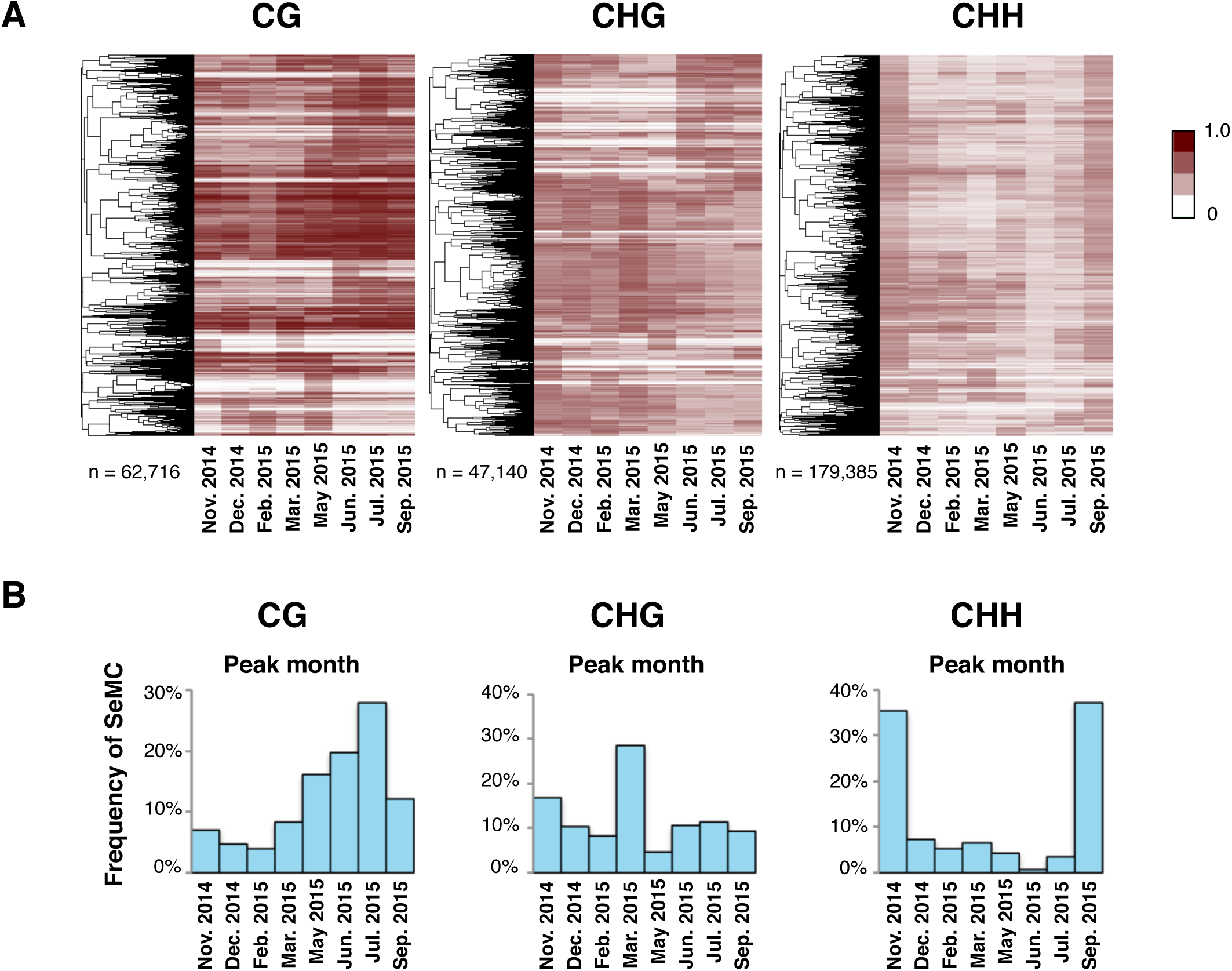
Annual patterns of seasonally-methylated cytosines (SeMCs). (*A*) Heatmaps of seasonally methylated cytosines (SeMCs) in CG, CHG, and CHH contexts (from left to right; n = 62,716, 47,140, 179,385, respectively). Each row indicates a series of DNA methylation ratios across a year in each position in the genome (0: unmethylated, 1; fully methylated). The dendrogram on the left of each heatmap represents the result of hierarchical clustering of SeMCs. Distance in the dendrogram was based on Pearson’s correlation coefficient. (*B*) Barplot showing the distribution of peak timings of methylation level for the SeMCs in CG, CHG, and CHH contexts (from left to right, respectively).

### CHH SeMCs in repetitive sequences

The distribution of SeMCs in the genome differed between the contexts of DNA methylation. The majority of SeMCs in CHH context (CHH-SeMCs) were found in intergenic regions (those annotated as neither exons nor introns) while CG-SeMCs and CHG-SeMCs were in both genic and intergenic regions (Fig. 4A). Given that large fractions of the eukaryotic genome are intergenic regions and consist of repetitive sequences such as transposable elements (26, 27), we compared the level of CHH methylation for all repetitive sequences in the genome of *A. halleri*. The median level of methylation was the highest in autumn, i.e., September and November (Fig. 4*B*), and the pattern was similar to that of whole-genome CHH-SeMCs. An example of seasonal changes in CHH methylation level in a LINE/L1 showed low and high levels in February-March and September-November, respectively (Fig. 4*C*). The highest methylation levels in autumn were detected in other selected families of repetitive sequences, such as LTR/Copia, LTR/Gypsy, LINE/L1, DNA/MULE-MuDR, DNA/hAT, and RC/Helitron (Fig. S3).

**Fig. 4.**
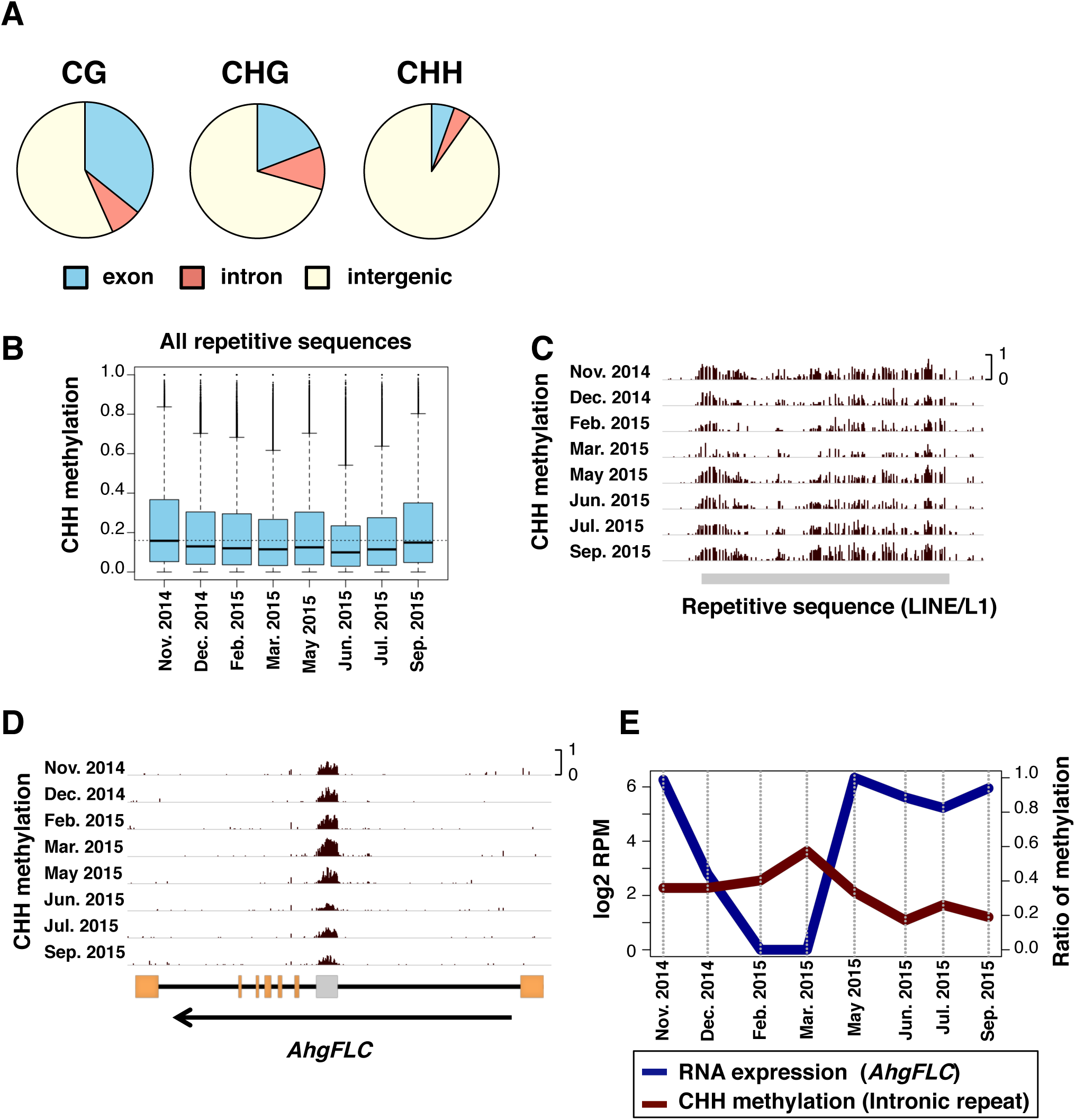
SeMCs in the CG, CHG, and CHH contexts, and seasonal patterns in CHH DNA methylation levels. (*A*) Pie charts indicating locations (exon, intron, and intergenic regions) of seasonally methylated cytosines (SeMCs) in CG, CHG, and CHH contexts (from left to right, respectively). (*B*) Boxplots of CHH methylation at 8 time points in repetitive elements. The boxes span from the first to the third quartiles, the thick black bars inside the boxes are the medians, whiskers above and below the boxes represent 1.5 × interquartile ranges from the quartiles. Dotted line indicates the median CHH methylation level in Nov. 2014. (*C* and *D*) Browser views for seasonal patterns of CHH methylation on repeat sequences in one of LINE/L1 sequences (*C*) and *AhgFLC* locus (*D*). Orange rectangles indicates exons. A gray rectangle in the *AhgFLC* locus indicates a repetitive sequence. (*E*) Comparison between RNA expression of *AhgFLC* and CHH methylation of the repetitive sequence in its intron.

In addition, we found some repetitive sequences that showed a unique seasonal pattern of CHH methylation. For example, in the intronic repeat at the locus of *FLOWERING LOCUS C* homolog (*AhgFLC*). *AhgFLC* expression is regulated seasonally and suppresses flowering when it is upregulated (20). We found that a SINE-like repetitive sequence in the first intron of *AhgFLC* showed a seasonal change of CHH methylation. The level was the highest in March and the lowest in June (Fig. 4*D*). Interestingly, this seasonal pattern was opposite to that of RNA expression of *AhgFLC* (Fig. 4*E*). The presence of interactions between transcription and CHH methylation in the intragenic repeat at *AhgFLC* is expected.

### Gene-body CG methylation (gbM) and constant RNA expression

Previous studies have reported that gbM is localized to active genes in a wide range of eukaryotes (5, 6). In *A. thaliana*, DNA methylation at the gene body is primarily observed in CG context (2, 3, 25). We examined seasonal patterns of gbM in *A. halleri* and found that the level of gbM was constant across seasons, both in medians and quantiles of all genes (Fig. S4*A*), and at the individual gene level (Fig. S4*B*).

In order to examine the potential role of gbM in gene regulation, we tested whether the level of gbM associated to seasonal patterns of RNA expression using previously published data of a two-year seasonal RNA-seq of *A. hallei* at the same study site (21). We found that genes with high gbM often have constant levels of RNA expression across the year. One such example was *AhgPP2AA3* (Fig 5 *A* and *B*), the homolog of *PROTEIN PHOSPHATASE 2A SUBUNIT A3*, a gene that is known as one that is constantly-expressed under various conditions in *A. thaliana* (28). This is in contrast to the situation of *AhgFLC* locus, a representative of seasonally expressed genes, which lacks CG methylation from the entire locus, except for the repetitive sequence in the first intron (Fig. 5 *C* and *D*). Based on these observations, we hypothesized that gbM would be associated to constant RNA expression across seasons. To test this hypothesis, we calculated the seasonal average and range of RNA expression for each gene, then compared them between genes with different levels of gbM (Fig. 5 *E* and *F*). Genes that were highly methylated in CG context showed relatively high average levels of RNA expression (Fig. 5*E*). The magnitude of seasonal changes in RNA expression of these genes decreased with increasing levels of gbM (Fig 5*F*). These results are consistent with the hypothesis mentioned above. The genes with the highest methylation level (in ‘group 5’ in Fig 5 *E* and *F*) were weakly enriched with four GOs of basic functions (Table S3).

**Fig. 5.**
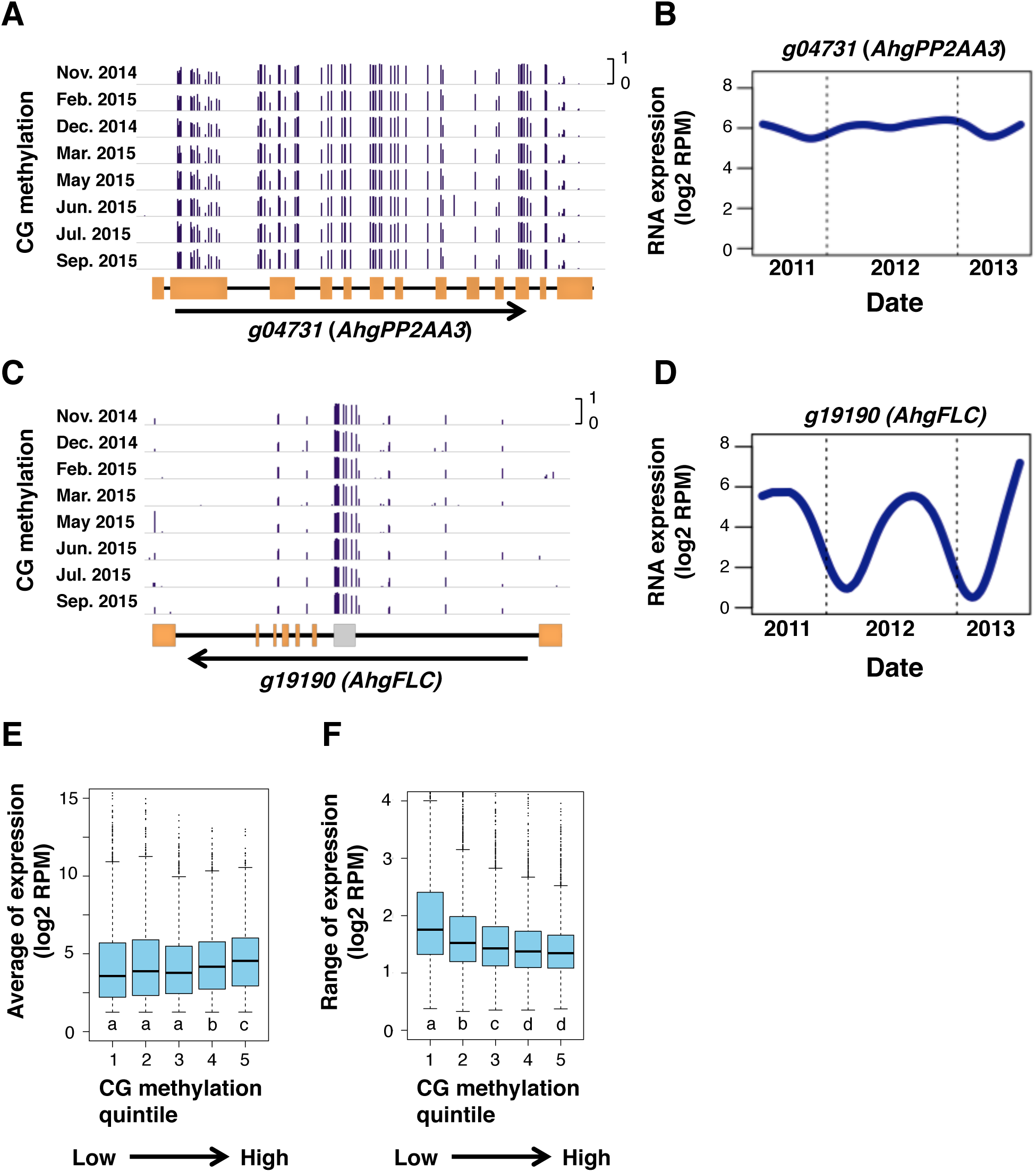
CG DNA methylation and stability of gene expression. (*A*-*D*) Browser views of seasonal patterns of CG methylation (*A* and *C*) and the two-year dynamics of RNA expression level (*B* and *D*; from September 2011 to August 2013) for *AhgPP2AA3* (*g04731*) and *AhgFLC* (*g19190*), respectively. Orange rectangles indicate exons. A gray rectangle indicates a repetitive sequence in the *AhgFLC* locus. (E, F) Boxplots showing relationship between DNA methylation in CG context and the average (E) and range (F) of RNA expression of genes. Only expressed genes (average expression level (log2(RPM) > 1)) are used for these analyses. Genes were split into five bins according to quintiles of genic DNA methylation in CG context. The boxes span from the first to the third quartiles, the bands inside the boxes are the medians, whiskers above and below the boxes represent 1.5 × interquartile ranges from the quartiles. Different letters represent significant differences between groups in the Mann-Whitney test, *P* < 0.01 adjusted for multiple comparisons.

## Discussion

In this study, we examined the seasonal patterns of DNA methylation at CG, CHG, and CHH contexts in *Arabidopsis halleri,* under natural conditions. We identified the genomic sites that showed seasonal changes in DNA methylation. Our observations suggest that seasonal factors in the natural habitat could affect DNA methylation differently according to its context and location in the genome. We would like to note here that, since our data set came from a single clonal individual for a single year, the genetic and non-genetic effects between different clones should be explored in future studies by applying WGBS to a population-level study with multiple repeat samples.

CHH methylation showed seasonal changes in diverse repetitive elements, and the level of DNA methylation was high in autumn, and low in winter. In *A. thaliana*, it has been reported that the level of CHH methylation in transposable elements was higher in 16°C environments relative to 10°C (13). Additionally, it has been suggested that RNA-directed DNA methylation (RdDM) pathway, which is required for maintenance and *de novo* methylation and mainly targets transposable elements, are involved in heat tolerance (29). This suggests that our observation on CHH SeMCs could reflect the temperature-dependence of the regulation of CHH methylation.

Seasonal changes detected in CHH methylation at a repetitive element in *AhgFLC* was an interesting example showing how genic heterochromatin behaves in genes. We observed that the seasonal pattern of CHH methylation in this particular repetitive element was opposite to that of the expression of *AhgFLC* gene. *FLC* and its homologs in Brassicaceae species contain conserved structures in its second intron called VRE (vernalization response element), which is involved in responsiveness to cold treatment (30, 31). Expression of *FLC* has been reported to be repressed by long-term cold treatment (vernalization), accompanied by enrichment of tri-methylation of histone H3 lysine 27 (H3K27me3) at the TSS site in the beginning of cold treatment, and then by H3K27me3 accumulation across gene body region after the plants returned to warmer temperatures (32). On the contrary, the expression of genes is associated with the removal of H3K27me3 from their bodies (33). These reports and our results suggest that epigenetic regulation of *AhgFLC* gene might be responsible for seasonal removal of CHH methylation in its intronic repeat. The disruption of transcription in genes has been reported to induce ectopic CHH methylation in *A. thaliana* (34).

Large-scale patterns of DNA methylation were constant throughout the year for all three contexts, although there were seasonal changes in DNA methylation at some cytosine sites. Constant levels of CG methylation were observed in gene body, and, furthermore, we found that genes with high levels of gbM showed seasonal stability in their gene expression. Although the function of gbM is still unclear (35), our results support previous reports, in *A. thaliana*, that gbM associates with genes modestly expressed among different tissues and experimental conditions (6, 36-38), and imply the importance of gbM under seasonal environment in a natural habitat.

Currently, we cannot entirely explain the patterns of DNA methylation in SeMCs, particularly in CG or CHG contexts. It is very likely that there are unidentified processes associated with environmental responses of DNA methylation that have not been studied under laboratory conditions. As mentioned above, the question remains of how the epigenetic variations in plants have been established in natural fluctuating environment (39, 40). Here, we focused on representative environmental effects on DNA methylation in a single clonal individual. In future studies of DNA methylation, we need to evaluate not only genotype effects, but also, so called ‘linage effects’ that represent past genetic and environmental effects. Furthermore, we should test to what extent seasonal changes in DNA methylation contribute to the variation in phenotypes in natural environments.

## Materials and Methods

### Plant materials

This study was conducted in a natural population of *Arabidopsis halleri* subsp. *gemmifera* located in central Japan (Omoide-gawa site, Naka-ku, Taka-cho, Hyogo Pref., 35°06’ N, 134°55’ E, altitude 190–230 m). Details of the study site have been described previously (19, 20). Leaf samples were collected at noon on the following dates: November 11 and December 22, 2014, and February 9, March 24, May 7, June 23, July 28, and September 8, 2015. In the study site, *A. halleri* forms patches of rosettes that consist of clonally propagated plants and sometimes genetically-related seed-originated plants. Originally six small patches of plants were chosen for leaf sampling, three of them were used for further analyses because the others were heavily damaged by deer herbivory between November 11 and December 22 in 2014. At each sampling date, we harvested multiple mature and intact leaves from each plant. Each leaf was ca. 3-cm long, and weighed ca. 0.1 g. To detect DNA methylation and RNA expression in the same set of leaves, a small piece was collected from each leaf for RNA extraction. For DNA extraction, leaves were frozen in an ethanol bath with dry ice, then stored at −80 °C. For RNA extraction, the small pieces of leaves were stored in RNA*later* solution (Invitrogen, Carlsbad, CA, USA), then stored at −20 °C according to the manufacturer’s instructions. The data for one patch were analyzed and shown in the main text, because the other two patches were found to be genetically mixed (Fig. S5). The samples analyzed here were confirmed to share whole-genome SNPs at levels that were safely judged to be a single clone (Fig. S5).

### DNA extraction and WGBS library preparation

Genomic DNA was extracted from collected leaves (two leaves per plant, ca. 0.2 g) using the CTAB method (41). Libraries for WGBS were constructed as described previously (42). Sequencing was performed with the Illumina Hiseq 2500 system.

### Processing of WGBS data

Sequenced reads were trimmed using Trimmomatic (43). Trimmed reads were mapped onto the genome sequence of *A. halleri* (44) using Bismark (45) and Bowtie2 (46). Repetitive sequences in the genome were detected using RepeatModeler (47), and repeats with at least 50 bp length were used for further analyses. The level of DNA methylation was calculated for each context using the ratio of the number of methylated cytosines to the number of total sequenced cytosine included in any region of the genome. Efficient bisulfite treatment (> 99% in all samples) was confirmed using the level of DNA methylation in unmethylated lambda DNA (Table S4). SeMC was defined as any cytosine differently methylated between at least two time points, detected by Fisher’s exact test, with genome-wide FDRs that were calculated using Storey’s method (48). To draw the heatmaps of methylation of SeMCs, cluster 3.0 (49) and Java Treeview (50) were used. To draw a circos plot for scaffold-wide DNA methylation, Circos software (51) was used. To make browser views of DNA methylation, Integrated Genome Viewer (52) was used. To draw a dendrogram of collected samples, MethylExtract (53), VCF-kit (https://vcf-kit.readthedocs.io), and Dendroscope3 (54) were used.

### Transcriptome analysis

RNA extraction and library preparation for RNA-seq were performed using the shotgun type method of BrAD-seq protocol (55). Sequencing was performed with the Illumina Hiseq 2500 system. Mapping of the reads and calculation of RPM were processed as described previously (21) except that the reference sequence was replaced with a newer annotation (44). To calculate seasonal average and range of expression of genes more precisely, weekly sampled RNA-seq data from a two-year period (21) were re-analyzed. To quantify the expression, Kallisto software was used (56). Calculation of seasonal average and range of RNA expression was based on previously described methods (21).

## Data availability

Raw WGBS and RNA-Seq reads are available under the DDBJ BioProject PRJDB7785.

## Acknowledgement

We thank Jiro Sugisaka for technical assistance, Takaomi Hatakeyama for information on *A. halleri* genome sequence, Atsushi J. Nagano for advice on analysis of RNA-seq data, and Kentaro K. Shimizu for comments on the manuscript. Computations were partially performed on the NIG supercomputer at NIG, Japan. This study was supported by JSPS Grant-in-Aid for Scientific Research (S) 26221106 and JST CREST JPMJCR15O1 to HK; Grant-in-Aid for JSPS Fellows by Japanese Society for Promoting Science (17J02659) to TI; Systems Functional Genetics Project of the Transdisciplinary Research Integration Center, ROIS, Japan to A.T., A.F., Y.T., and T.K.

## Author contributions

H.K. and T.K. conceived the study. T.I., H.N., H.K., and T.K. designed the experiments. H.N. collected samples. Y.T., A.T., and A.F. performed whole-genome bisulfite sequencing. N.E. and M.N.H. performed RNA sequencing. T.I. analyzed the data. T.I. and H.K. wrote the paper and incorporated comments from co-authors.

---

**Fig. S1.** Genome-wide bulk DNA methylation level at eight time points across a year. Genome-wide bulk DNA methylation levels are shown in CG, CHG, and CHH context for the whole genome (*A*), and gene and repetitive sequences (*B*). Eight sampling timings are represented by the different shades.

**Fig. S2.** DNA methylation was seasonally stable at a majority of CG, CHG, and CHH sites. Histograms of seasonal differences of DNA methylation (max. – min.) for all cytosine sites (*A*), and for SeMCs (*B*) in CG, CHG, and CHH contexts.

**Fig. S3.** Seasonal patterns in CHH DNA methylation in repetitive elements that belong to the six major families of transposable elements (TEs). The boxes span from the first to the third quartiles, the thick black bars inside the boxes are the medians, whiskers above and below the boxes represent 1.5 × interquartile ranges from the quartiles.

**Fig. S4.** Seasonal patterns in CG gene body methylation (gbM) across a year. (*A*) Boxplot of DNA methylation in genes in CG context at eight time points across a year. The boxes span from the first to the third quartiles, the thick black bars inside the boxes are the medians, whiskers above and below the boxes represent 1.5 × interquartile ranges from the quartiles. (*B*) A histogram of seasonal difference for DNA methylation in genes (max. – min.) in CG context.

**Fig. S5.** Some patches of A. halleri turned to be genetically mixed. A dendrogram shows Kimura’s genetic distance using genome-wide SNPs among the samples in three patches for eight timepoints. The samples from replicate 1 were judged to be a single clone. A. halleri is an obligate outcrossing species with a self-incompatible breeding system, and therefore a large portion of SNPs are expected to be heterozygous. Because heterozygous SNPs can be designated as homozygous to either of the alleles in a certain probability, a terminal radial branching is expected to be observed even for genetically identical plants that belong to a single clonal patch (Rep. 1).

## References

1. Law JA, Jacobsen SE (2010) Establishing, maintaining and modifying DNA methylation patterns in plants and animals. Nat Rev Genet 11:204–220.

2. Cokus SJ, et al. (2008) Shotgun bisulphite sequencing of the *Arabidopsis* genome reveals DNA methylation patterning. Nature 452:215–219.

3. Lister R, et al. (2008) Highly integrated single-base resolution maps of the epigenome in *Arabidopsis*. Cell 133:523–536.

4. Lister R, et al. (2009) Human DNA methylomes at base resolution show widespread epigenomic differences. Nature 462:315–322.

5. Feng S, et al. (2010) Conservation and divergence of methylation patterning in plants and animals. Proc Natl Acad Sci USA 107:8689–8694.

6. Zemach A, McDaniel IE, Silva P, Zilberman D (2010) Genome-wide evolutionary analysis of eukaryotic DNA methylation. Science 328:916–919.

7. Huff JT, Zilberman D (2014) Dnmt1-Independent CG methylation contributes to nucleosome positioning in diverse eukaryotes. Cell 156:1286–1297.

8. Niederhuth CE, et al. (2016) Widespread natural variation of DNA methylation within angiosperms. Genome Biol 17:194.

9. Cubas P, Vincent C, Coen E (1999) An epigenetic mutation responsible for natural variation in floral symmetry. Nature 401:157–161.

10. Manning K, et al. (2006) A naturally occurring epigenetic mutation in a gene encoding an SBP-box transcription factor inhibits tomato fruit ripening. Nat Genet 38:948–952.

11. Ong-Abdullah M, et al. (2015) Loss of Karma transposon methylation underlies the mantled somaclonal variant of oil palm. Nature 525:533–537.

12. Dowen RH, et al. (2012) Widespread dynamic DNA methylation in response to biotic stress. Proc Natl Acad Sci USA 109:E2183–E2191.

13. Dubin MJ, et al. (2015) DNA methylation in *Arabidopsis* has a genetic basis and shows evidence of local adaptation. eLife 4:elife.05255.

14. Secco D, et al. (2015) Stress induced gene expression drives transient DNA methylation changes at adjacent repetitive elements. eLife 4:e09343.

15. Becker C, et al. (2011) Spontaneous epigenetic variation in the *Arabidopsis thaliana* methylome. Nature 480:245–249.

16. Schmitz RJ, et al. (2011) Transgenerational epigenetic instability is a source of novel methylation variants. Science 334:369–373.

17. Shimizu KK, Kudoh H, Kobayashi MJ (2011) Plant sexual reproduction during climate change: gene function *in natura* studied by ecological and evolutionary systems biology. Ann Bot 108:777–787.

18. Kudoh H (2016) Molecular phenology in plants: *in natura* systems biology for the comprehensive understanding of seasonal responses under natural environments. New Phytol 210:399–412.

19. Kudoh H, Honjo MN, Nishio H, Sugisaka J (2018) The long-term “*in natura*” study sites of *Arabidopsis halleri* for plant transcription and epigenetic modification analyses in natural environments. Methods Mol Biol 1830:41–57.

20. Aikawa S, Kobayashi MJ, Satake A, Shimizu KK, Kudoh H (2010) Robust control of the seasonal expression of the Arabidopsis *FLC* gene in a fluctuating environment. Proc Natl Acad Sci USA 107:11632–11637.

21. Nagano AJ, et al. (2019) Annual transcriptome dynamics in natural environments reveals plant seasonal adaptation. Nat Plants 5:74–83.

22. Stroud H, et al. (2013) Non-CG methylation patterns shape the epigenetic landscape in *Arabidopsis*. Nat Struct Mol Biol 21:64–72.

23. Kawashima T, Berger F (2014) Epigenetic reprogramming in plant sexual reproduction. Nat Rev Genet 15:613–624.

24. He Y, Ecker JR (2015) Non-CG methylation in the human genome. Annu Rev Genomics Hum Genet 16:55–77.

25. Seymour DK, Koenig D, Hagmann J, Becker C, Weigel D (2014) Evolution of DNA methylation patterns in the Brassicaceae is driven by differences in genome organization. PLoS Genet 10:e1004785.

26. Fedoroff NV (2012) Transposable elements, epigenetics, and genome evolution. Science 338:758–767.

27. Tenaillon MI, Hollister JD, Gaut BS (2010) A triptych of the evolution of plant transposable elements. Trends Plant Sci 15:471–478.

28. Czechowski T (2005) Genome-wide identification and testing of superior reference genes for transcript normalization in Arabidopsis. Plant Physiol 139:5–17.

29. Popova OV, Dinh HQ, Aufsatz W, Jonak C (2013) The RdDM Pathway Is required for basal heat tolerance in *Arabidopsis*. Mol Plant 6:396–410.

30. Sung S, et al. (2006) Epigenetic maintenance of the vernalized state in *Arabidopsis thaliana* requires LIKE HETEROCHROMATIN PROTEIN 1. Nat Genet 38:706–710.

31. Castaings L, et al. (2014) Evolutionary conservation of cold-induced antisense RNAs of *FLOWERING LOCUS C* in *Arabidopsis thaliana* perennial relatives. Nat Commun 5:4457. doi:10.1038/ncomms5457.

32. Song J, Irwin J, Dean C (2013) Remembering the prolonged cold of winter. Curr Biol 23:R807–R811.

33. Buzas DM, Robertson M, Finnegan EJ, Helliwell CA (2011) Transcription-dependence of histone H3 lysine 27 trimethylation at the Arabidopsis polycomb target gene *FLC*. Plant J 65:872–881.

34. Inagaki S, et al. (2010) Autocatalytic differentiation of epigenetic modifications within the Arabidopsis genome. EMBO J 29:3496–3506.

35. Zilberman D (2017) An evolutionary case for functional gene body methylation in plants and animals. Genome Biol 18:87.

36. Zhang X, et al. (2006) Genome-wide high-resolution mapping and functional analysis of DNA methylation in *Arabidopsis*. Cell 126:1189–1201.

37. Aceituno FF, Moseyko N, Rhee SY, Gutiérrez RA (2008) The rules of gene expression in plants: Organ identity and gene body methylation are key factors for regulation of gene expression in *Arabidopsis thaliana*. BMC Genomics 9:438.

38. Coleman-Derr D, Zilberman D (2012) Deposition of histone variant H2A.Z within gene bodies regulates responsive genes. PLoS Genet 8:e1002988.

39. Richards EJ (2006) Inherited epigenetic variation — revisiting soft inheritance. Nat Rev Genet 7:395–401.

40. Weigel D, Colot V (2012) Epialleles in plant evolution. Genome Biol 13:249.

41. Saghai-Maroof MA, Soliman KM, Jorgensen RA, Allard RW (1984) Ribosomal DNA spacer-length polymorphisms in barley: mendelian inheritance, chromosomal location, and population dynamics. Proc Natl Acad Sci USA 81:8014–8018.

42. Fu Y, et al. (2013) Mobilization of a plant transposon by expression of the transposon-encoded anti-silencing factor. EMBO J 32:2407–2417.

43. Bolger AM, Lohse M, Usadel B (2014) Trimmomatic: a flexible trimmer for Illumina sequence data. Bioinformatics 30:2114–2120.

44. Briskine RV, et al. (2016) Genome assembly and annotation of *Arabidopsis halleri*, a model for heavy metal hyperaccumulation and evolutionary ecology. Mol Ecol Resour 17:1025–1036.

45. Krueger F, Andrews SR (2011) Bismark: a flexible aligner and methylation caller for Bisulfite-Seq applications. Bioinformatics 27:1571–1572.

46. Langmead B, Salzberg SL (2012) Fast gapped-read alignment with Bowtie 2. Nat Methods 9:357–359.

47. Smit AFA, Hubley R (2008-2015) RepeatModeler Open-1.0. http://www.repeatmasker.org.

48. Storey JD (2002) A direct approach to false discovery rates. J R Stat Soc Series B Stat Methodol 64:479–498.

49. de Hoon MJL, Imoto S, Nolan J, Miyano S (2004) Open source clustering software. Bioinformatics 20:1453–1454.

50. Saldanha AJ (2004) Java Treeview—extensible visualization of microarray data. Bioinformatics 20:3246–3248.

51. Krzywinski M, et al. (2009) Circos: An information aesthetic for comparative genomics. Genome Res 19:1639–1645.

52. Robinson JT, et al. (2011) Integrative genomics viewer. Nat Biotechnol 29:24–26.

53. Barturen G, Rueda A, Oliver JL, Hackenberg M (2013) Methylextract. doi:10.5281/zenodo.7144.

54. Huson DH, Scornavacca C (2012) Dendroscope 3: An Interactive Tool for Rooted Phylogenetic Trees and Networks. Syst Biol 61:1061–1067.

55. Townsley BT, Covington MF, Ichihashi Y, Zumstein K, Sinha NR (2015) BrAD-seq: Breath Adapter Directional sequencing: a streamlined, ultra-simple and fast library preparation protocol for strand specific mRNA library construction. Front Plant Sci 6. doi:10.3389/fpls.2015.00366.

56. Bray NL, Pimentel H, Melsted P, Pachter L (2016) Near-optimal probabilistic RNA-seq quantification. Nat Biotechnol 34:525–527.

