## Supplemental Figures and Tables for "Seasonal stability and dynamics of DNA methylation in plants in a natural environment"

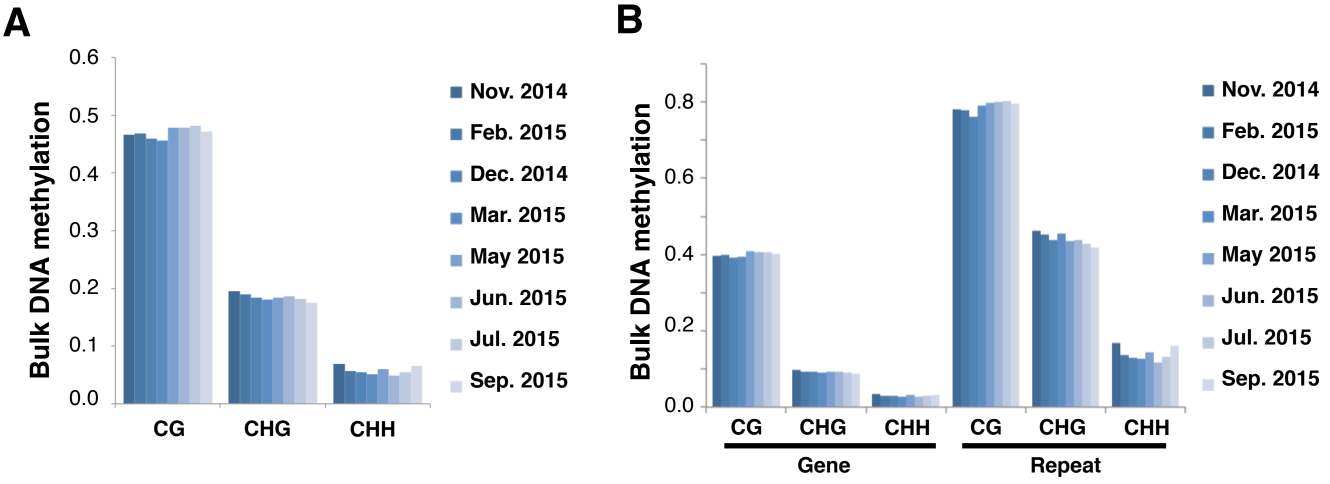


**Fig. S1. Genome-wide bulk DNA methylation level at eight time points across a year.** Genome-wide bulk DNA methylation levels are shown in CG, CHG, and CHH context for the whole genome (*A*), and gene and repetitive sequences (*B*). Eight sampling timings are represented by the different shades.

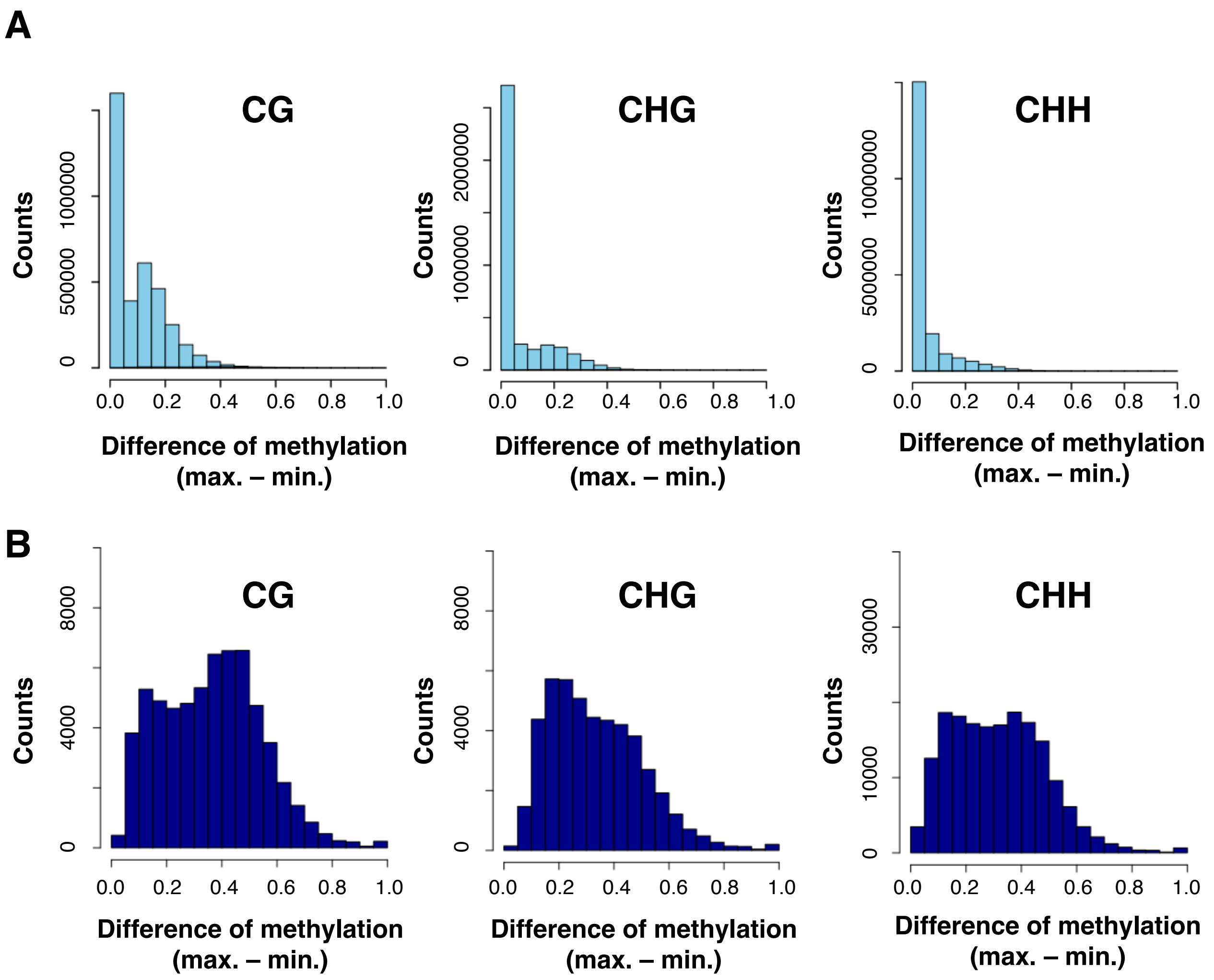


**Fig. S2. DNA methylation was seasonally stable at a majority of CG, CHG, and CHH sites.** Histograms of seasonal differences of DNA methylation (max. – min.) for all cytosine sites (*A*), and for SeMCs (*B*) in CG, CHG, and CHH contexts.

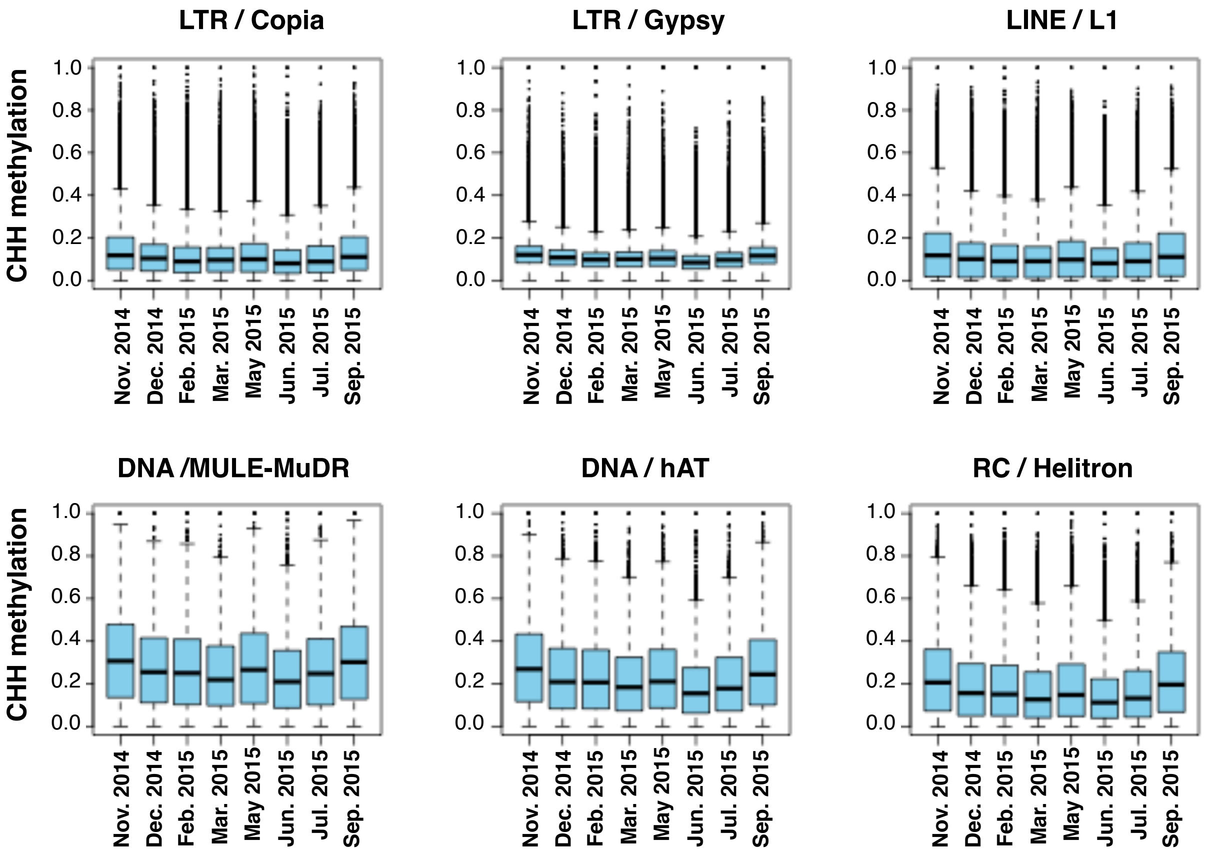


**Fig. S3. Seasonal patterns in CHH DNA methylation in repetitive elements that belong to the six major families of transposable elements (TEs).** The boxes span from the first to the third quartiles, the thick black bars inside the boxes are the medians, whiskers above and below the boxes represent 1.5 × interquartile ranges from the quartiles.

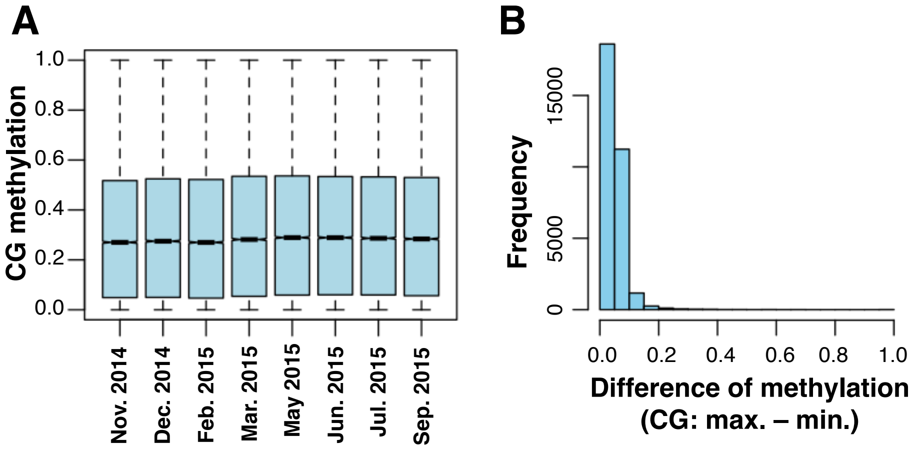


**Fig. S4. Seasonal patterns in CG gene body methylation (gbM) across a year.** (*A*) Boxplot of DNA methylation in genes in CG context at eight time points across a year. The boxes span from the first to the third quartiles, the thick black bars inside the boxes are the medians, whiskers above and below the boxes represent 1.5 × interquartile ranges from the quartiles. (*B*) A histogram of seasonal difference for DNA methylation in genes (max. – min.) in CG context.

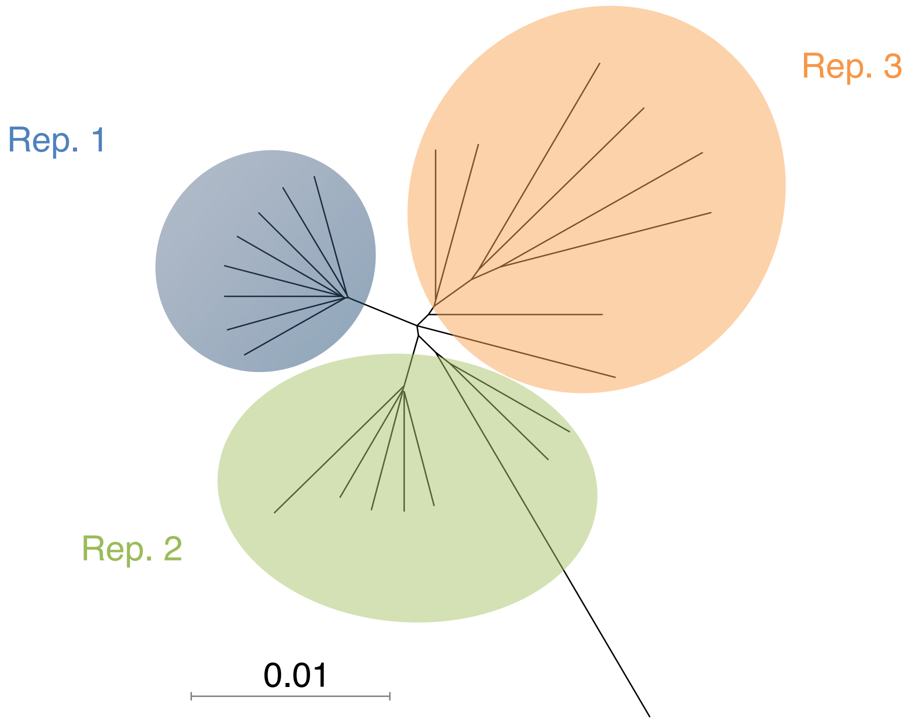


**Fig. S5. Some patches of A. halleri turned to be genetically mixed.** A dendrogram shows Kimura's genetic distance using genome-wide SNPs among the samples in three patches for eight timepoints. The samples from replicate 1 were judged to be a single clone. A. halleri is an obligate outcrossing species with a self-incompatible breeding system, and therefore a large portion of SNPs are expected to be heterozygous. Because heterozygous SNPs can be designated as homozygous to either of the alleles in a certain probability, a terminal radial branching is expected to be observed even for genetically identical plants that belong to a single clonal patch (Rep. 1).

Table S1. Summary of sequencing and mapping for the genome-wide DNA methylation data at 8 sampling times.

| Sampling time | Number of processed reads | Number of mapped reads | Mapping efficiency |
|---|---|---|---|
| Nov. 2014 | 90,520,893 | 40,941,176 | 45.20% |
| Dec. 2014 | 87,303,234 | 37,824,643 | 43.30% |
| Feb. 2015 | 92,067,366 | 38,579,387 | 41.90% |
| Mar. 2015 | 90,115,863 | 34,016,251 | 37.70% |
| May. 2015 | 91,491,555 | 38,257,975 | 41.80% |
| Jun. 2015 | 89,141,892 | 37,461,447 | 42.00% |
| Jul. 2015 | 91,279,854 | 39,459,871 | 43.20% |
| Sep. 2015 | 89,745,835 | 40,284,890 | 44.90% |

Table S2. Summary of correlation between DNA methylation and density of repeats in 100 kbp windows at 8 sampling times.

| | Pearson's Correlation coefficient | | |
|---|---|---|---|
| Sampling time | CG | CHG | CHH |
| Nov. 2014 | 0.64 | 0.68 | 0.70 |
| Dec. 2014 | 0.63 | 0.67 | 0.68 |
| Feb. 2015 | 0.63 | 0.68 | 0.69 |
| Mar. 2015 | 0.63 | 0.67 | 0.68 |
| May. 2015 | 0.64 | 0.67 | 0.68 |
| Jun. 2015 | 0.64 | 0.67 | 0.67 |
| Jul. 2015 | 0.64 | 0.68 | 0.68 |
| Sep. 2015 | 0.64 | 0.68 | 0.70 |

Table S3. Enrichment of Gene Ontology in the 'group 5' of Fig. 5 E and F.

Ontology: Process

| GO term | Description | P-value | FDR q-value |
|---|---|---|---|
| GO:0043161 | proteasome-mediated ubiquitin-dependent protein catabolic process | 9.04E-04 | 1.00E+00 |

Ontology: Function

| GO term | Description | P-value | FDR q-value |
|---|---|---|---|
| GO:0051011 | microtubule minus-end binding | 2.69E-04 | 2.15E-01 |

Ontology: Component

| GO term | Description | P-value | FDR q-value |
|---|---|---|---|
| GO:0099568 | cytoplasmic region | 1.94E-04 | 9.29E-02 |
| GO:0005938 | cell cortex | 1.94E-04 | 4.64E-02 |

Table S4. Efficiency of bisulfite treatment in this study.

| Sampling time | Efficiency |
|---|---|
| Nov. 2014 | 99.34% |
| Dec. 2014 | 99.38% |
| Feb. 2015 | 99.42% |
| Mar. 2015 | 99.39% |
| May. 2015 | 99.29% |
| Jun. 2015 | 99.42% |
| Jul. 2015 | 99.48% |
| Sep. 2015 | 99.46% |
